# Quantifying subclinical and longitudinal microvascular changes following episcleral plaque brachytherapy (EPB) using spectral-domain OCT angiography

**DOI:** 10.1101/860148

**Authors:** Kyle M. Green, Brian C. Toy, Bright S. Ashimatey, Debarshi Mustafi, Richard L. Jennelle, Melvin A. Astrahan, Zhongdi Chu, Ruikang K. Wang, Jonathan Kim, Jesse L. Berry, Amir H. Kashani

**Author notes:** contributed equally (Kyle M. Green, Brian C. Toy). **Meeting Presentation:** American Society of Retina Specialists Annual Meeting, Vancouver, British Columbia, July 22, 2018. **Communicating Author:** Amir H. Kashani MD PhD, Institute for Biomedical Therapeutics, 1450 San Pablo St., 6^th^ Floor, Los Angeles, CA 90033.

## Abstract

**Background:** I-125 episcleral plaque brachytherapy (EPB) is standard-of-care for globe-conserving treatment of medium-sized choroidal melanomas. Radiation retinopathy is a potential consequence of treatment, characterized by deleterious effects on retinal microvasculature. We investigated the application of Optical Coherence Tomography Angiography (OCTA) for detecting and longitudinally monitoring I-125 episcleral plaque brachytherapy induced radiation retinopathy.

**Methods:** High resolution OCTA of the central 3×3mm macula were obtained from I-25 episcleral plaque brachytherapy treated and untreated fellow eyes of 62 patients. Capillary density (vessel skeleton density, VSD) and caliber (vessel diameter index, VDI) were quantified using previously validated semi-automated algorithms. Nonperfusion was also quantified as flow impairment regions (FIR). Exams from treated and fellow eyes obtained pre-treatment and at 6-month, 1-year, and 2-year intervals were compared using generalized estimating equation linear models. Dosimetry maps were used to evaluate spatial correlation between radiation dose and microvascular metrics.

**Results:** Mean time from treatment to last follow-up was 10.8 months. Mean±SD and median radiation dose at the fovea were 64.5 ± 76 Gy and 32.0 Gy, respectively. Preoperative logMAR (Snellen) mean visual acuity was 0.26 ± 0.05 (∼20/35) and 0.08 ± 0.02 (∼20/25) in treated and fellow eyes, respectively. At 6 months, treated eyes had significantly lower VSD (0.147 ± 0.003 vs 0.155 ± 0.002; *p* = 0.023) and higher FIR (1.95 ± 0.176 vs 1.45 ± 0.099; *p* = 0.018) compared to fellow eyes. There was a significant decrease in VSD and a corresponding increase in FIR even for treated eyes without clinically identifiable retinopathy at 6 months. VDI was significantly higher in treated eyes than in fellow eyes at 2 years (2.93 ± 0.022 vs 2.84 ± 0.016; *p* = 0.002). Microvascular changes were spatially correlated with a radiation gradient of 85-250 Gy across the fovea.

**Conclusions:** OCTA can be used to quantify and monitor EPB induced radiation, and can detect vascular abnormalities even in the absence of clinically observable retinopathy. OCTA may therefore be useful in investigating treatment interventions that aim to delay EPB-induced radiation retinopathy.

## Introduction

The development of radiation retinopathy (RR) following treatment of choroidal melanoma with episcleral plaque brachytherapy (EPB) can have deleterious effects on retinal microvasculature that leads to permanent visual decline. The Collaborative Ocular Melanoma Study (COMS) validated EPB as standard-of-care for globe-conserving treatment of medium-sized choroidal melanomas.(1) Despite the selection of iodine-125 (I-125) and gold shielding to minimize adverse radiation effects, visual morbidity remains high, with only 43% of patients maintaining visual acuity of 20/200 or better 3 years after treatment with standard COMS plaques.(2) Some reports indicate that adverse radiation effects can be partially mitigated through the use of 3D conformal treatment planning software and customized collimated plaques to decrease the radiation dose to critical visual structures (e.g. the fovea).(3, 4) The onset of RR varies greatly, ranging from as early as 1 month to 15 years, but it most commonly occurs between 6 months and 3 years after treatment.(5) The location of the tumor, and thus the dose to the fovea, is critical but not the sole risk factor. RR is primarily a vascular disease and shares many clinical features with diabetic retinopathy including damage to capillaries, which leads to variable degrees of hyperpermeability, retinal ischemia, and neovascularization.(6)

While clinical features of RR, including dot-blot hemorrhages, microaneurysms, and macular edema can be seen on exam, as with diabetic retinopathy, there is underlying damage present before these clinical features manifest. Fluorescein angiography (FA) can reveal early areas of non-perfusion and vascular leakage.(7-9) OCT angiography (OCTA) has been used to non-invasively demonstrate morphologic features of microvasculature with excellent resolution.(10-13) By generating detailed depth-resolved images, OCTA can potentially be used to detect and monitor capillary-level aberrations in blood flow at multiple timepoints early in the course of RR. To date, there have been a few studies that employ OCTA to assess microvascular changes in RR. Two recent studies used binarized OCT angiograms to demonstrate a decrease in parafoveal (14, 15) and peripapillary capillary (16, 17) density in irradiated eyes compared to fellow eyes. To our knowledge, no studies have performed longitudinal analysis to identify early microvascular changes (prior to 1 year) in treated eyes, nor have any used OCTA to explore a possible spatial correlation between these changes and radiation dose.

In the present study, we employed longitudinal analysis of OCT angiograms to further determine what quantifiable morphologic differences exist in the microvasculature between treated and fellow eyes over time, as well as how early in the course of RR these differences can be detected. More specifically, we used previously described OCTA metrics, (11, 18) vessel skeleton density (VSD) and vessel diameter index (VDI), to quantitatively assess changes in retinal vascular networks. We have previously employed these metrics to quantify vascular density and diameter in diabetic retinopathy and uveitis.(11, 13) We also report flow impairment region (FIR), which is further detailed in the methods section, to quantify areas of subclinical non-perfusion larger than a set threshold.

## Methods

Approval for this study was obtained from the Institutional Review Board of the University of Southern California, and the described research adhered to the tenets of the Declaration of Helsinki. This was a retrospective, consecutive series of 62 adult patients treated with I-125 episcleral plaque brachytherapy (EPB) for medium sized choroidal melanoma by one of two ocular oncologists (JB, JK) at the USC Roski Eye Institute. Any subject with history of intraocular melanoma and plaque brachytherapy was eligible for inclusion. Treatment planning and surgery were performed as previously described with stereotactic plaque brachytherapy radiation treatment planning platform (Plaque Simulator; Eye Physics, LLC; Los Alamitos, CA); a dose of 85 Gy at a rate of 0.5 Gy/hr was prescribed to the apex of the lesion or to 5mm height, whichever was greater with a 2mm margin at the base.(19) Subjects with media opacity impairing visualization of the macula, pre-existing retinal vascular disease (diabetic retinopathy, retinal vein occlusion, choroidal neovascularization), or pre-existing subretinal fluid or macular edema prior to plaque placement were excluded from the study. Subjects with direct tumor involvement in the 3×3mm perifoveal region were also excluded as this was the area assessed by OCTA. Patient demographics, including age and gender, were abstracted from the medical record. Clinical data collected included visual acuity, radiation dose to fovea, follow-up time, and presence of clinically evident radiation retinopathy as determined by the Finger criteria.(8) Visual acuity was reported as logMAR (Snellen equivalent).

### OCTA Imaging and Image Analysis

OCTA was performed in both the irradiated eye and the fellow non-irradiated eye for each patient during the patient’s regularly scheduled clinic visits. Imaging was performed with a commercially available, FDA-approved, spectral-domain OCTA platform (Angioplex; Zeiss Meditec; Dublin, CA). High resolution 3×3mm OCT angiograms centered on the fovea with a signal strength greater than 7/10 were included for analysis. Images with significant motion artifacts that obscured the vascular architecture were excluded from analysis. For any eye with multiple images on a single date, the highest quality image was chosen. A previously described semi-automated algorithm was used for quantitative analysis of vessel skeletal density (VSD), vessel diameter index (VDI), and flow impairment region (FIR).(11, 18) Briefly, VSD is defined as a unitless proportion of the total length (in pixels) of all skeletonized vessels divided by the total number of pixels in the image window, which reflects capillary density. VDI is defined as a unitless proportion of the total vessel area divided by the total skeletonized vessel area, which reflects average vessel diameter. Lastly, FIR is defined as the sum of avascular areas in an image larger than a pre-defined threshold area, which in this study was set at 0.002mm^2^ to reflect the maximum threshold for physiologic intercapillary distance. This value was based on an estimate from histologic analysis that the avascular periarteriolar region is ∼50μm.(20) A 0.002mm^2^ threshold closely approximates the area of a circle with this diameter.

### Data Analyses

The effect of radiation therapy on the OCTA metrics (VSD, FIR and VDI) was assessed longitudinally. Data acquired included pretreatment exams obtained prior to surgery, as well as post-treatment exams binned into 6-month (range 3-9 months), 1-year (range 9-18 months) and 2-year (range 18-30 months) groups. Summary OCTA metrics were compared between irradiated and fellow eyes at the different time intervals using generalized estimating equation (GEE) linear models. In the GEE models, the OCTA metrics were each used as predictor variables of the treatment status of the eyes—irradiated eye versus fellow eye. Summary OCTA metrics for treated eyes were also compared between baseline and the various follow-up timepoints. Statistical significance was defined when the p-value associated with the odds ratio of the univariate model was less than 0.05. GEE models allow for the analysis of longitudinal repeated measures, as well as correlated fellow eye data.(21) When the number of radiated and fellow eyes were balanced, paired t-statistic or Wilcoxon sign-ranked tests were also used.

The radiation dose-related changes of the OCTA metrics were also investigated. The OCTA metrics between high-dose eyes (foveal radiation >45 Gy) were compared to low-dose eyes (foveal radiation <15 Gy). These thresholds were chosen based on published dose tolerance limits of the retina.(22) A second exploratory approach was adapted to assess if there was spatial correlation between radiation dose and microvascular density within the 3×3mm foveal regions imaged. To evaluate this “within eye” correlation, the last acquired OCT angiograms (over the defined study period) of the irradiated eyes were investigated and five eyes which displayed spatial gradients in microvascular density were subjectively selected for further evaluation. EPB dosimetry maps of these eyes were then generated using Eye Physics Plaque Simulator software (Eye Physics, LLC; Los Alamitos, CA) developed previously at the University of Southern California.(23-25) For each case, dosimetry maps were superimposed on both the original OCT angiograms and their corresponding fundus photos for analysis.

## Results

We report the results of 62 participants who underwent EPB therapy. Table 1 summarizes the demographic and clinical characteristics of the study population. Table 2 summarizes the results of the OCTA metrics compared between EPB-treated and untreated fellow eyes.

**Table 1.**
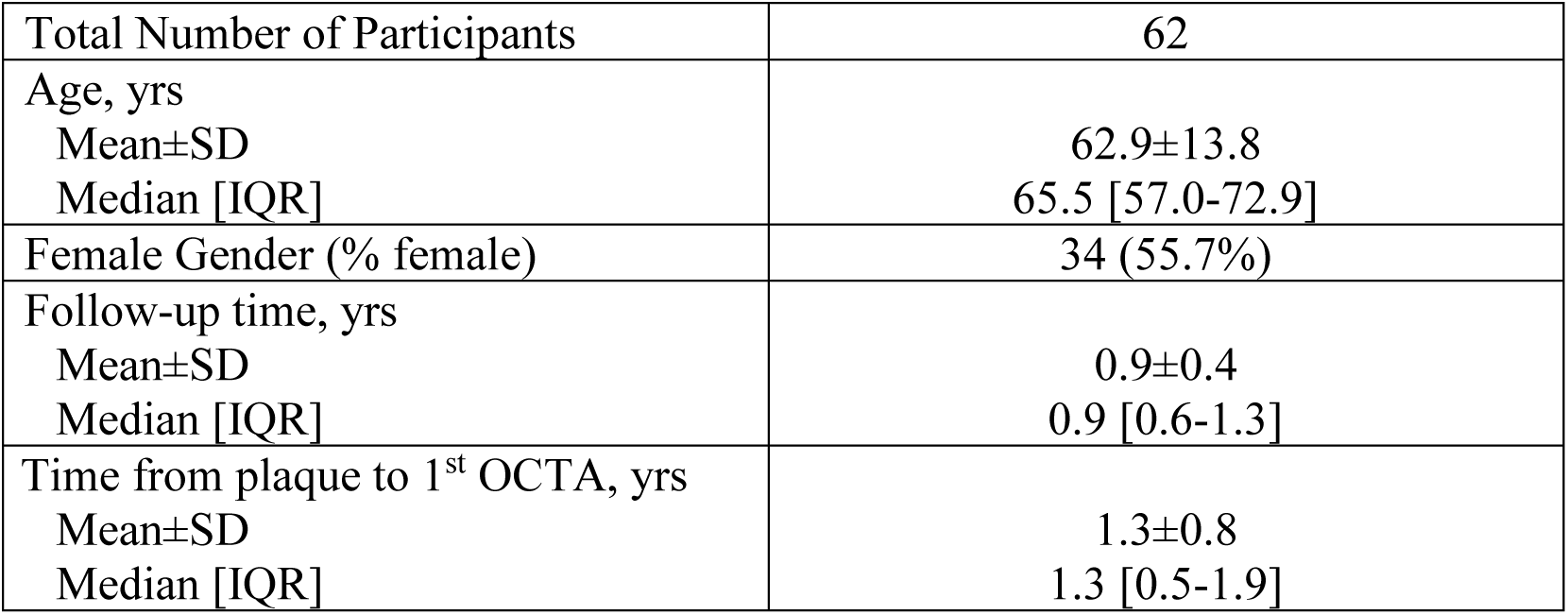
Patient demographics and clinical data.

**Table 2.**
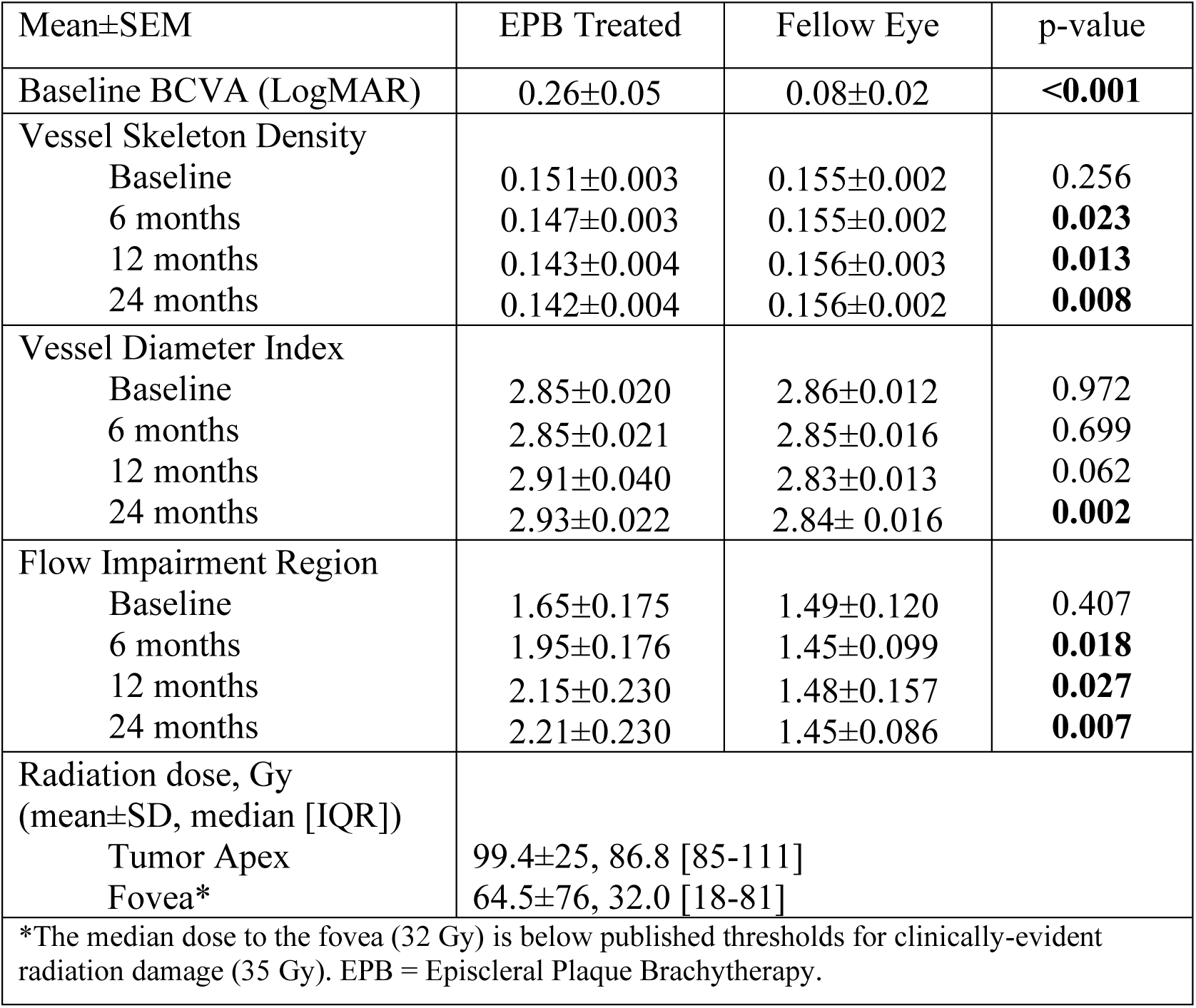
Clinical and OCTA measures by eye.

### Baseline

Prior to EPB, eyes with melanoma had significantly lower visual acuity compared to fellow eyes; however, there were no significant differences in VSD, VDI, or FIR at baseline between eyes with melanoma and the contralateral eyes (Table 2).

### Six Month Follow-Up

Fifteen subjects had OCT angiograms at 6 months after EPB. Only one of these (representing 6.7%) demonstrated even minimal evidence of radiation retinopathy on clinical examination. However, the VSD and FIR metrics of OCTA assessment showed significantly lower VSD and higher FIR for the treated eyes compared to fellow eyes respectively (Table 2). These changes can also be appreciated qualitatively in maps of VSD and FIR (Figure 1). Importantly, among treated eyes that had no clinically identifiable radiation retinopathy at this follow-up period, and also had pre-treatment exams for direct comparison (n=5), there was still a significantly decreased VSD (0.146±0.011 [6 months] vs 0.158±0.005 [baseline]; *p* = 0.035) and an increased FIR (1.76±0.665 [6 months] vs 1.28±0.339 [baseline]; *p* = 0.043).

**Fig 1.**
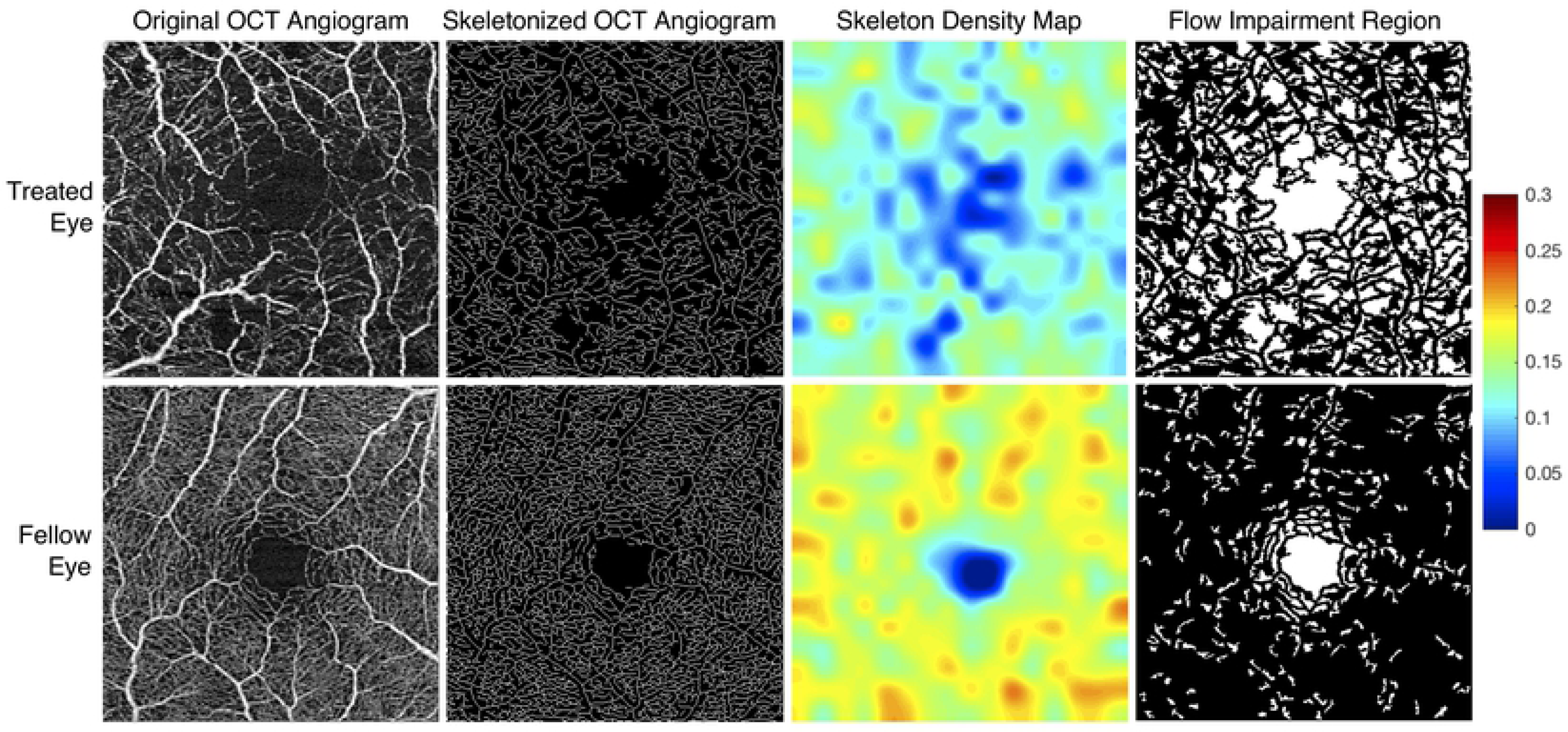
Processed OCT angiograms from treated and fellow eyes of a single patient. OCT angiograms from the treated (OS) and fellow eye (OD) of a 20-year-old female demonstrate marked qualitative differences in parafoveal vessel density (column 1). The OCT angiogram of each eye was obtained 263 days (8.6 months) following placement in the treated eye with a 46.0 Gy dose at the fovea. Visual acuity at the time of image acquisition was 20/350 in the treated eye and 20/20 in the fellow eye. Skeletonized OCT angiograms with accompanying skeleton density heat maps were generated (columns 2 and 3). Warmer colors reflect areas of greater vessel skeleton density (VSD), with relative differences defined on the accompanying color scale demonstrating decreased VSD in the treated eyes. Pseudocolor flow impairment maps (column 4) demonstrate absent flow signal (white areas). The flow impairment region was markedly increased in the treated eye.

### One Year Follow-Up

At 12 months after EPB, visual acuity was 0.29±0.075 (∼20/40) and 0.06±0.016 (∼20/20) in the treated and fellow eyes respectively (p = 0.005). 25% (4 of 16) of treated eyes with exams at this time point demonstrated at least minimal evidence of radiation retinopathy on clinical examination. Treated eyes also showed a significant lower VSD and higher FIR compared to fellow eyes (Table 2).

### Two Year Follow-Up

At 24 months after EPB, visual acuity was 0.37±0.09 (∼20/45) and 0.10±0.06 (∼20/25) in the treated and fellow eyes respectively (*p* = 0.015). 75% (12 of 16) of treated eyes with exams at this time point demonstrated at least minimal evidence of radiation retinopathy on clinical examination. Treated eyes also showed a significantly lower VSD compared to fellow eyes (Table 2). In general, the difference in all metrics between treated and fellow eyes grew over time and corresponded with increasing rates of clinically identifiable radiation retinopathy in treated eyes (Figure 2).

**Fig 2.**
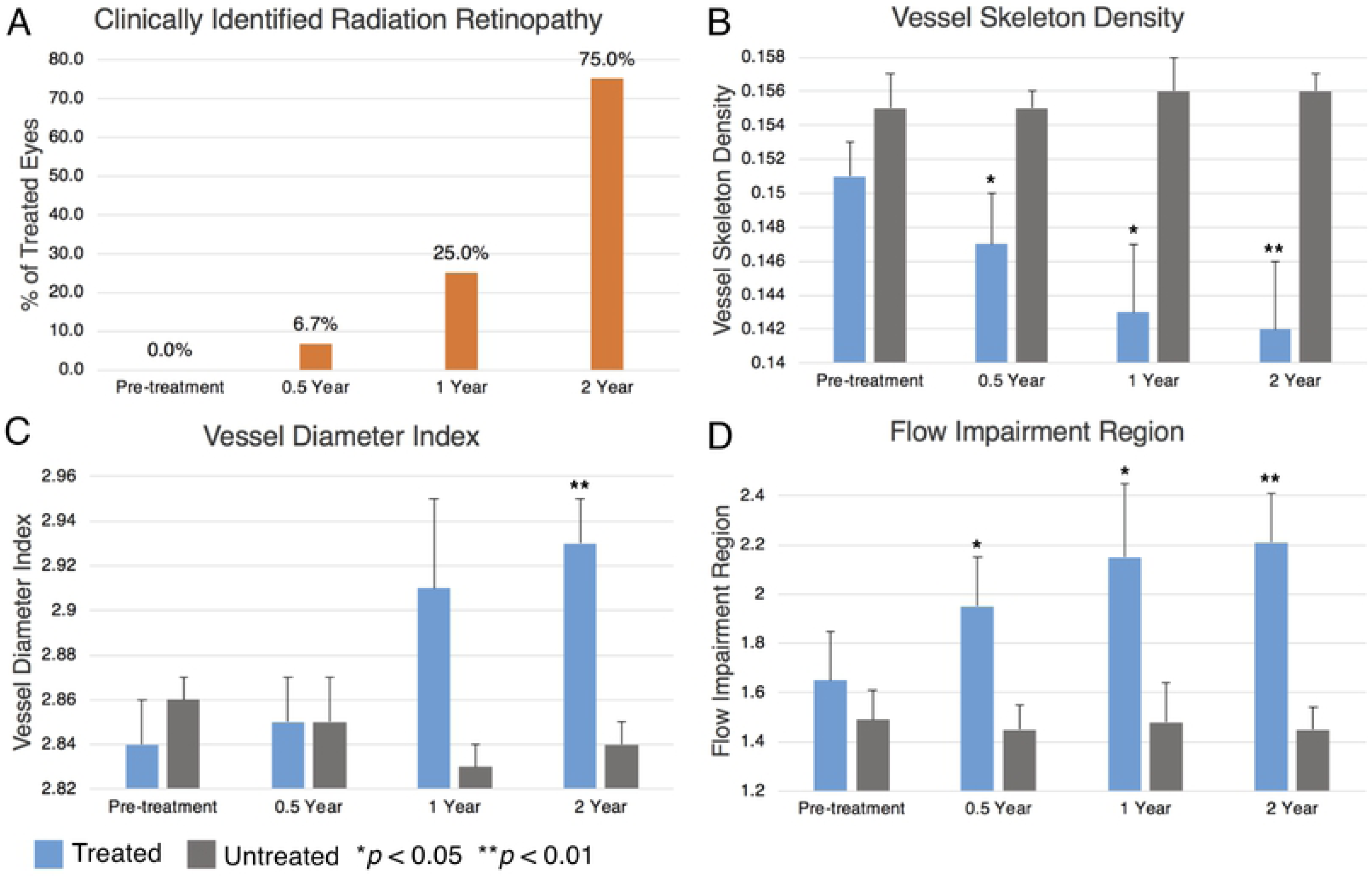
Longitudinal clinical and quantitative OCTA data. All panels reflect data from our overall cohort. Over the course of our 2-year follow-up period, there was an increasing percentage of treated eyes with clinically identifiable radiation retinopathy at each interval (A). Compared to fellow eyes over this period, treated eyes showed decreasing vessel skeleton density (VSD) (B), increasing flow impairment region (C), and increasing vessel diameter index (D). Relative significance between treated and fellow eyes at each time point is marked by asterisks, and error bars reflect standard error of the mean.

### Radiation Dose Correlation with OCTA Changes

We found significant differences in the OCTA metrics VSD and FIR over the follow-up period when the overall cohort was divided into high and low dose foveal radiation subgroups (>45 Gy [n=9] vs <15 Gy [n=3]): VSD (0.145±0.002 [high dose] (26) vs 0.154±0.001 (27), *p* < 0.0001) and FIR (2.04±0.10 (26) vs 1.59±0.06 (27), *p* < 0.0001). The VDI metric was however not significantly different between the high dose and low dose classification (2.88±0.02 (26) vs 2.83±0.08 (27), *p* = 0.21).

The five 3×3mm OCT angiograms selected for the “within eye” dose-effect analysis had the following range of radiation doses across the fovea: Case 1 - 85-250 Gy; Case 2 - 30-70 Gy; Case 3 - 25-60 Gy; Case 4 - 40-60 Gy; and Case 5 - 8-12 Gy. Of these, the case with the greatest radiation gradient across the fovea (Case 1) had EPB dosimetry gradient that spatially correlated with the microvascular gradient on the 3×3mm OCT angiogram. The longitudinal OCTA findings of Case 1 are illustrated in Figure 3, and the registered EPB dosimetry map and OCTA microvasculature is illustrated in Figure 4. The dose-dependent nature of impaired perfusion over time can be appreciated from Figure 3 when the EPB dosimetry map in Figure 4 is taken into account. The remaining cases did not appear to have any spatially correlated microvascular changes within the 3×3mm window.

**Fig 3.**
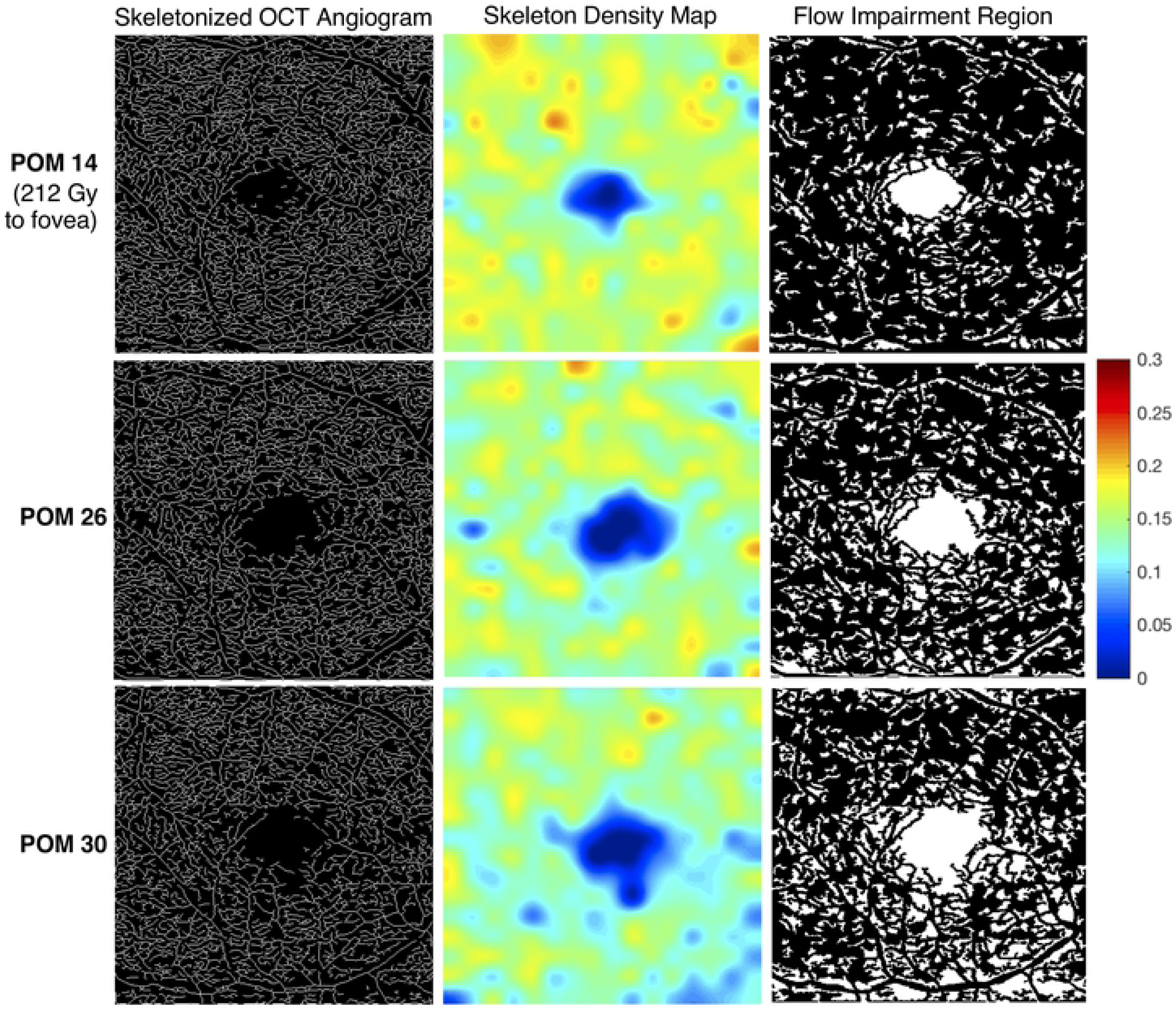
Longitudinal skeleton density and flow impairment maps of a treated eye. This patient is a 65-year-old male who received 212 Gy to the fovea (OD), with a range of 85-250 Gy across the standard 3×3mm OCT angiogram (Case 1). OCT angiograms were acquired at post-operative months (POM) 14, 26, and 30. The visual acuity of the treated eye at these dates was 20/25, 20/25, and 20/80 respectively. The visual acuity of the fellow eye at the same time points (OCTA images not shown) was 20/25, 20/20, and 20/25, respectively. In the skeletonized image, impaired perfusion is visible inferiorly at POM 26 compared to POM 14, with worsening perfusion at POM 30 (column 1). The loss of skeleton density is more clearly visualized in the heat map (column 2). Warmer colors reflect areas of greater vessel skeleton density, with relative differences defined on the color scale. A parallel trend is seen in the flow impairment region images (column 3).

**Fig 4.**
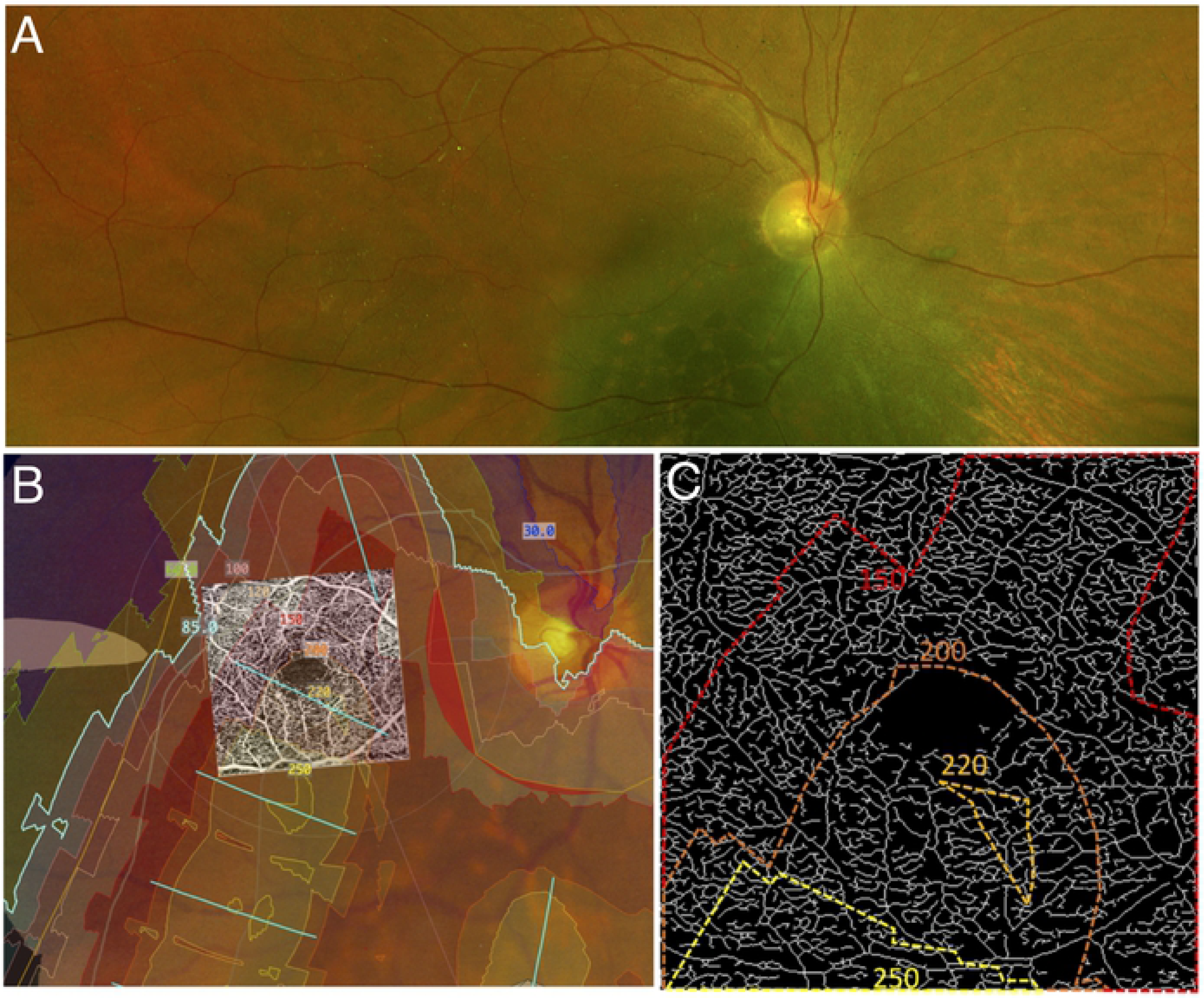
Spatial correlation of parafoveal microvascular changes with radiation dose. Panel A shows the pre-treatment fundus image of a subject (Case 1) showing the choroidal melanoma. Panel B is a computed dosimetry simulation projected onto the pre-treatment fundus image. A 3×3mm OCT angiogram of the eye was registered with the image using vessel bifurcation landmarks. Dosimetry contour lines and dosimetry tints delineate areas of the eye that received specific doses of radiation from the plaque. Panel C is an enlarged skeletonized 3×3mm OCT angiogram of the eye at post-op month 30 with the corresponding dosimetry contour lines. Note the inferior areas of decreased vascular density (impaired perfusion) in the 3×3mm image, which corresponds with the higher doses delivered inferiorly.

## Discussion

This study adds to a body of literature that has demonstrated retinal microvascular changes after episcleral plaque brachytherapy (EPB). Specifically, our study demonstrated a significant decrease in capillary density in EPB treated eyes earlier than previously reported and prior to clinically evident radiation retinopathy. It also demonstrated progressive decreases in density at intervals over a 2-year period. This was accomplished through the use of quantitative metrics that directly reflect microvascular density such as vessel skeleton density (VSD), and also indirectly such as flow impairment region (FIR). Significant changes in vessel diameter index (VDI) were also seen over this time period. In addition to these findings, we present a case with a large gradient (>165 Gy) of high-dose radiation across the fovea that appears to be spatially correlated to microvascular density. Collectively, these data suggest that capillary changes are occurring before clinically evident retinopathy, and that the magnitude of the radiation dose may correlate with the magnitude of the capillary damage in any given region. Furthermore, they highlight the potential utility of OCTA to monitor the progression of subtle changes in microvasculature over a period of months in treated eyes.

Our findings were consistent with those in prior studies that used OCTA to assess parafoveal vessel density in irradiated eyes. Say et al. and Cennamo et al quantified total vascular area using 3×3mm and 6×6mm binarized en-face images, respectively.(14, 15) Both demonstrated significant reduction in vessel area density in irradiated eyes compared to fellow eyes. Although the capillary densities in these previous studies were estimated as vessel area density, our preferred method for estimating capillary density is the skeletonized density (VSD). This is because VSD is not influenced by capillary morphologic changes such as vessel diameter, which may accompany vasculopathies, and is also minimally impacted by large caliber vessels. For brevity, our study only reports the VSD analysis as the measure for capillary density. Vessel diameter was approximated as an index -VDI - which we also demonstrate changes with worsening retinopathy. FIR, our third metric, complements VSD as an indirect measure of density and a direct measure of subclinical impaired perfusion. As FIR only accounts for avascular areas above a set threshold, it theoretically has a higher specificity (but lower sensitivity) for capillary dropout. For example, the loss of very minute areas of capillary flow may not result in an avascular area above our set threshold, and therefore would have no effect on FIR, but a definite effect on VSD.

The findings of our study highlight the potential use of OCTA for monitoring vascular changes in irradiated eyes over time. The vascular metrics can also serve as adjuncts to help grade the severity of radiation retinopathy. Several groups have aimed to develop effective grading schemes that use various imaging modalities, including ultra-wide field fluorescein angiography.(9) In 2005, Finger et al developed a system with four stages of severity that correlated with vision loss, based on a combination of findings from dye-based angiography and ophthalmoscopy.(8) Horgan et al later described in 2008 how OCT could be further added to identify macular edema, an early clinical feature of radiation retinopathy.(7) More recently, Veverka et al. suggested OCTA could also be used to help grade severity, demonstrating that it may detect RR prior to changes seen on OCT alone.(28)

Thus, we concur with the assertion that OCTA may be a powerful tool in determining the severity of radiation retinopathy, and also in detecting very early microvascular changes before the onset of retinopathy on exam. This has a wide variety of clinical applications. For example, OCTA can contribute relevant information for individualizing the time point for RR treatment intervention, and also provide sensitive biomarkers for comparing the efficacy of RR treatment regimens.(29-31) The use of various metrics as demonstrated in this study may noticeably increase the sensitivity of OCTA to capture early changes in RR, as subtle changes in density and vessel diameter are often challenging to appreciate qualitatively in the clinic setting. For clinical purposes, we suggest obtaining 3×3mm OCT angiograms in both eyes prior to EPB placement, and intermittently at follow-up visits for those with access to these devices. As our study has shown, significant microvascular changes can be seen within 6 months of treatment, suggesting repeat imaging may be prudent at early post-operative dates. Furthermore, as our understanding of the utility of OCTA continues to grow, longitudinal scans may prove useful in the long-term management of individuals with RR including indications for therapy.

Our exploration of a possible spatial correlation between radiation dose and capillary density was demonstrated in Case 1 (Figure 4) which had (by chance alone) a very steep gradient change for the radiation dose over the 3×3mm area of the macula which was imaged. Of note, Case 4 also showed a large area of ischemia nearest the high dose radiation in a wider 6×6mm window. The significantly lower resolution of 6×6mm OCTA scans precludes a detailed analysis of density changes in these scans. Our findings provide a basis for future studies assessing the within eye spatial relation between EPB dosimetry and microvasculature abnormalities to enhance the understanding of radiation dose on the retinal vasculature and the development of radiation retinopathy.

Some potential limitations of our study include those inherent to OCTA imaging, such as motion artifacts and floaters, which can interfere with efforts to accurately quantify vascular metrics. We aimed to control for this by excluding images with significant artifacts. Additionally, this study analyzed images from the 3×3mm OCTA scan pattern and may have missed some peripheral defects associated with EPB. However, larger scan patterns available at the time of this study did not have sufficient resolution to reliably detect capillary level changes, so use of the high resolution 3×3mm field was necessary. Future studies may consider images from 6×6mm or even larger windows if the resolution of the scans is sufficient. Furthermore, future studies may aim to generate dosimetry maps in a larger number of eyes, and employ more quantitative approaches to better evaluate the spatial relationship between EPB dosimetry and microvascular aberrations. Other limitations are from the retrospective nature of the data analyzed. For example, the images for the study were acquired during study visits which were determined on a case to case basis by the physician. Although we addressed the difference in the time intervals by binning, our findings can be refined by using a regularized and standardized time intervals across subjects.

In conclusion, we investigated OCTA changes associated with EPB treatment of choroidal melanoma and report significant changes in OCTA metrics at about 6 months or earlier, even when there were no clinically detectable signs of radiation retinopathy. The change in the OCTA metrics increased over time, and in a dose dependent manner. We infer that OCTA can be a viable tool for monitoring the effect of EPB on the retinal microvasculature and its findings may play a pivotal role in developing intervention modalities to delay or treat the occurrences of retinopathy after episcleral plaque brachytherapy.

## Abbreviations

EPB: Episcleral Plaque Brachytherapy
FA: Fluorescein Angiography
FIR: Flow Impairment Region
OCT(A): Optical Coherence Tomography (Angiography)
RR: Radiation Retinopathy
VDI: Vessel Diameter Index
VSD: Vessel Skeleton Density

